## Supplemental Figures for "Cis-Regulatory Hubs: a new 3D Model of Complex Disease Genetics with an Application to Schizophrenia"

### Supplementary

#### Figures

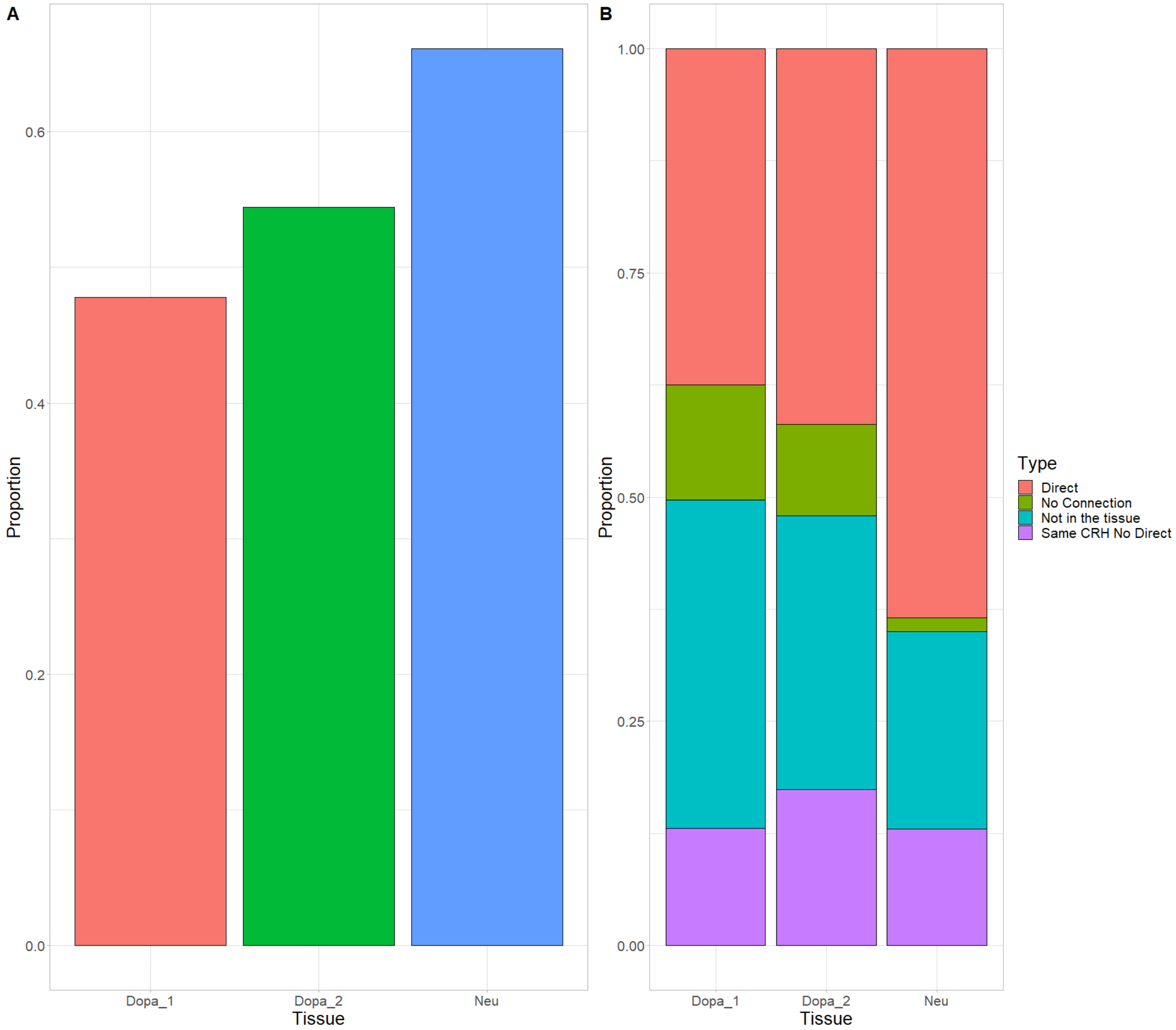


##### Figure S1

##### CRHs exhibit strong overlap between iPSC neurons and post-mortem brain tissues

**(A)** Proportion of distal elements observed in iPSC neurons found in the post-mortem brain tissues**. (B)** Proportion of pairs of promoter-distal element observed in iPSC-derived neurons strictly found in one of other post-mortem tissue (Direct), found in the tissue but not either as direct or indirect connection (No Connection), found in the tissue within the same CRH (Same CRH No Direct) or not found in the tissue. Results are reported relatively to the number of pairs observed in iPSC-derived neurons.

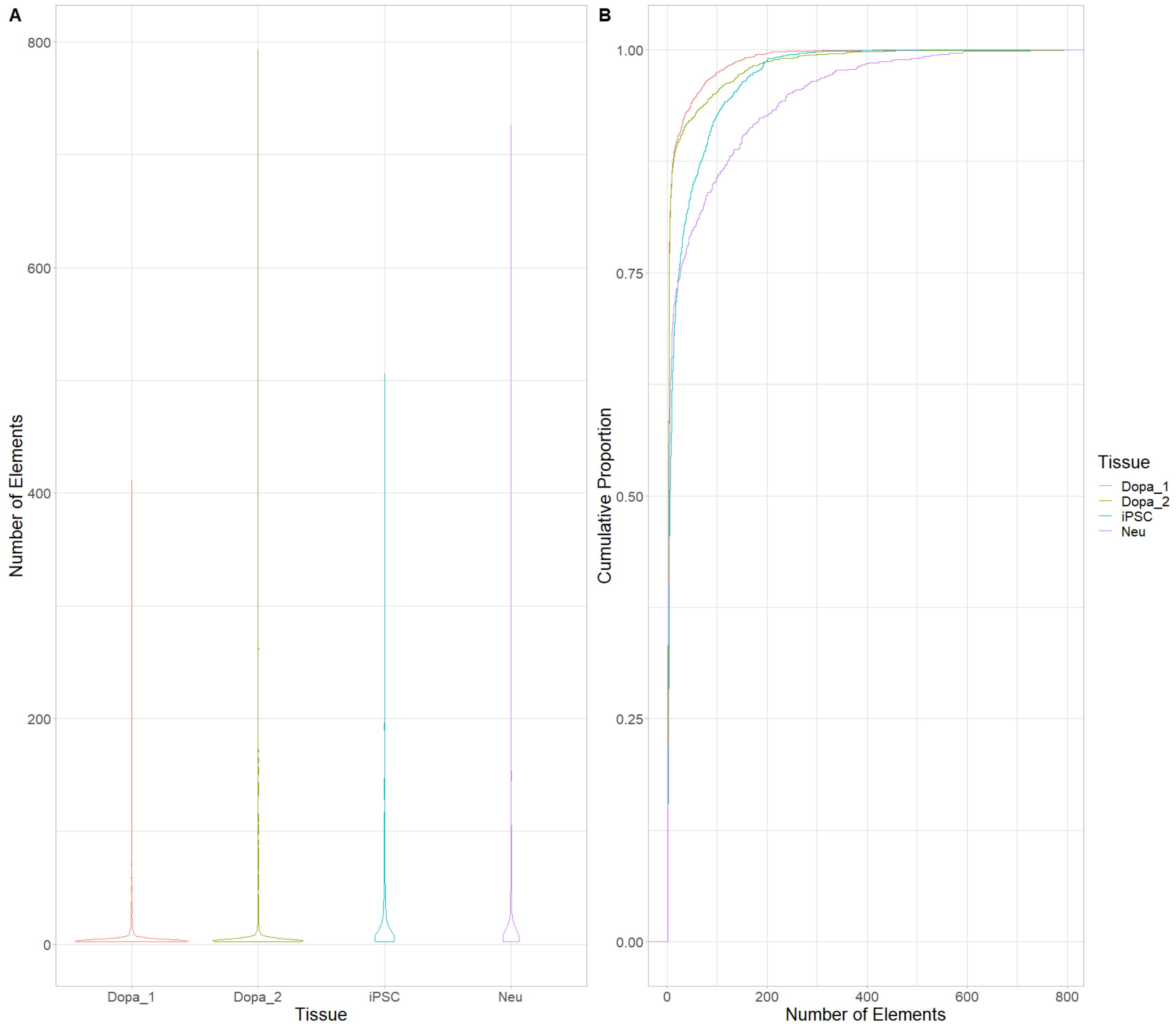


**Figure S2**

**CRHs in iPSC-derived neurons show "average behavior" compared to post-mortem tissues regarding number of elements**

**(A)** Violin plots of the number of elements included in CRHs by brain tissue. **(B)** Cumulative distribution of the number of elements within CRHs by tissue.

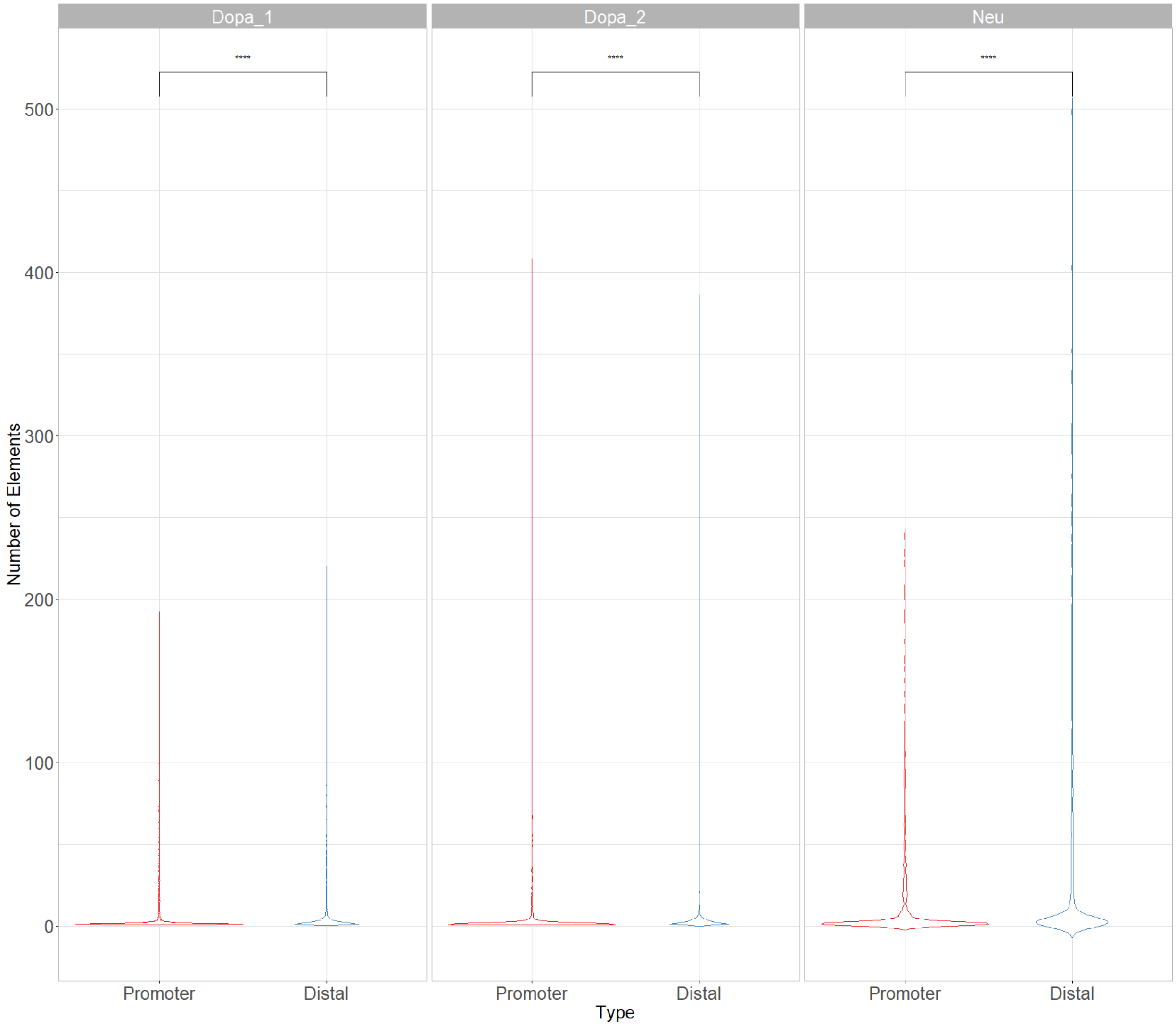


**Figure S3**

**In post-mortem tissues, CRHs are mainly composed by distal elements**

Violin plots of the number of promoters or distal elements across post-mortem brain tissues. Difference between number of elements were assessed with Wilcoxon signed-rank test.

Data Information: **** represent p-value <=0.0001.

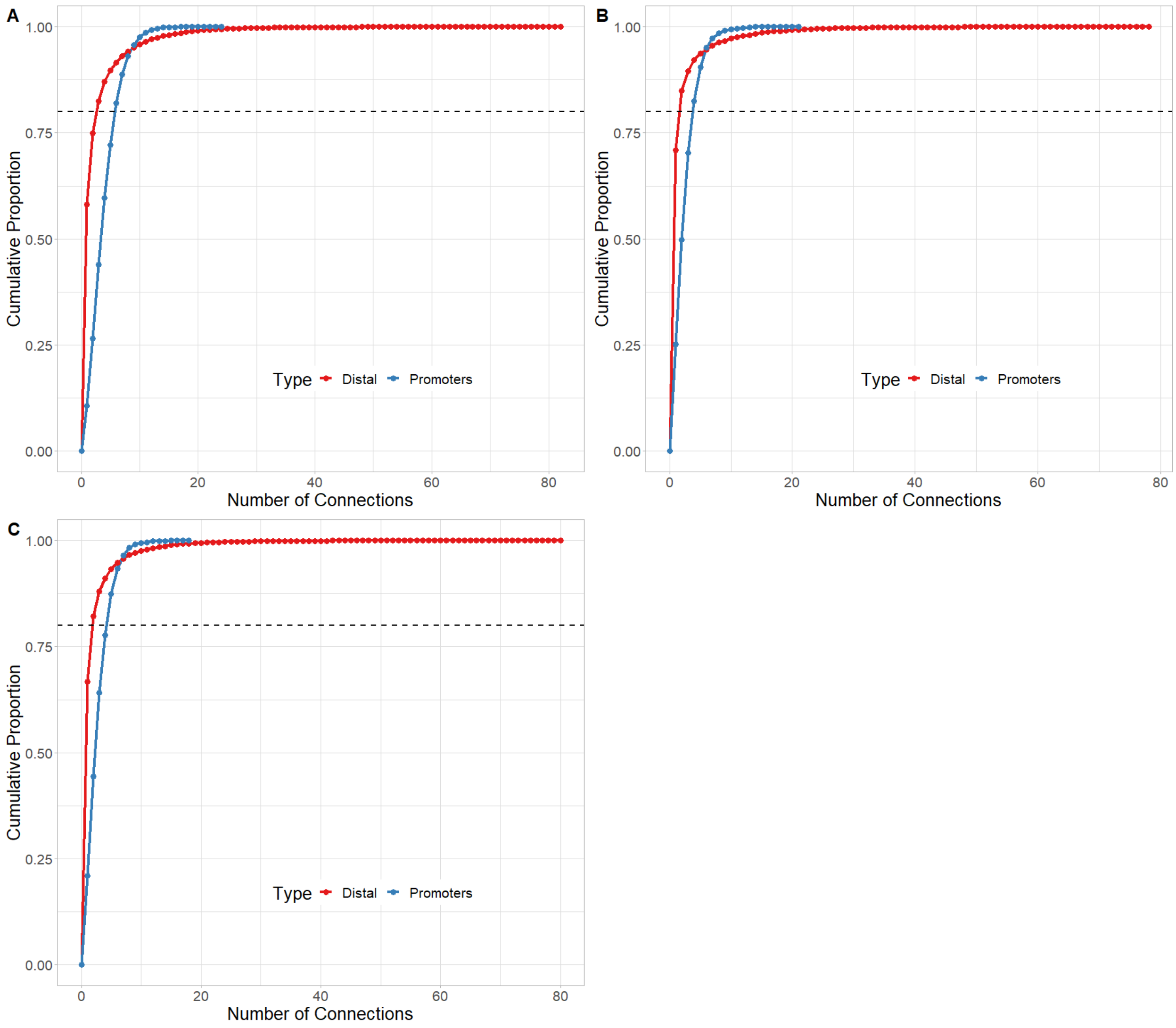


#### Figure S4

#### Promoters are more connected than distal elements across post-mortem tissues

Cumulative distribution function of the number of connections by promoter and distal element for **(A)** Neu, **(B)** Dopa_1, and **(C)** Dopa_2. Dotted line shows the number of connections where 80% are less or equal to this value.

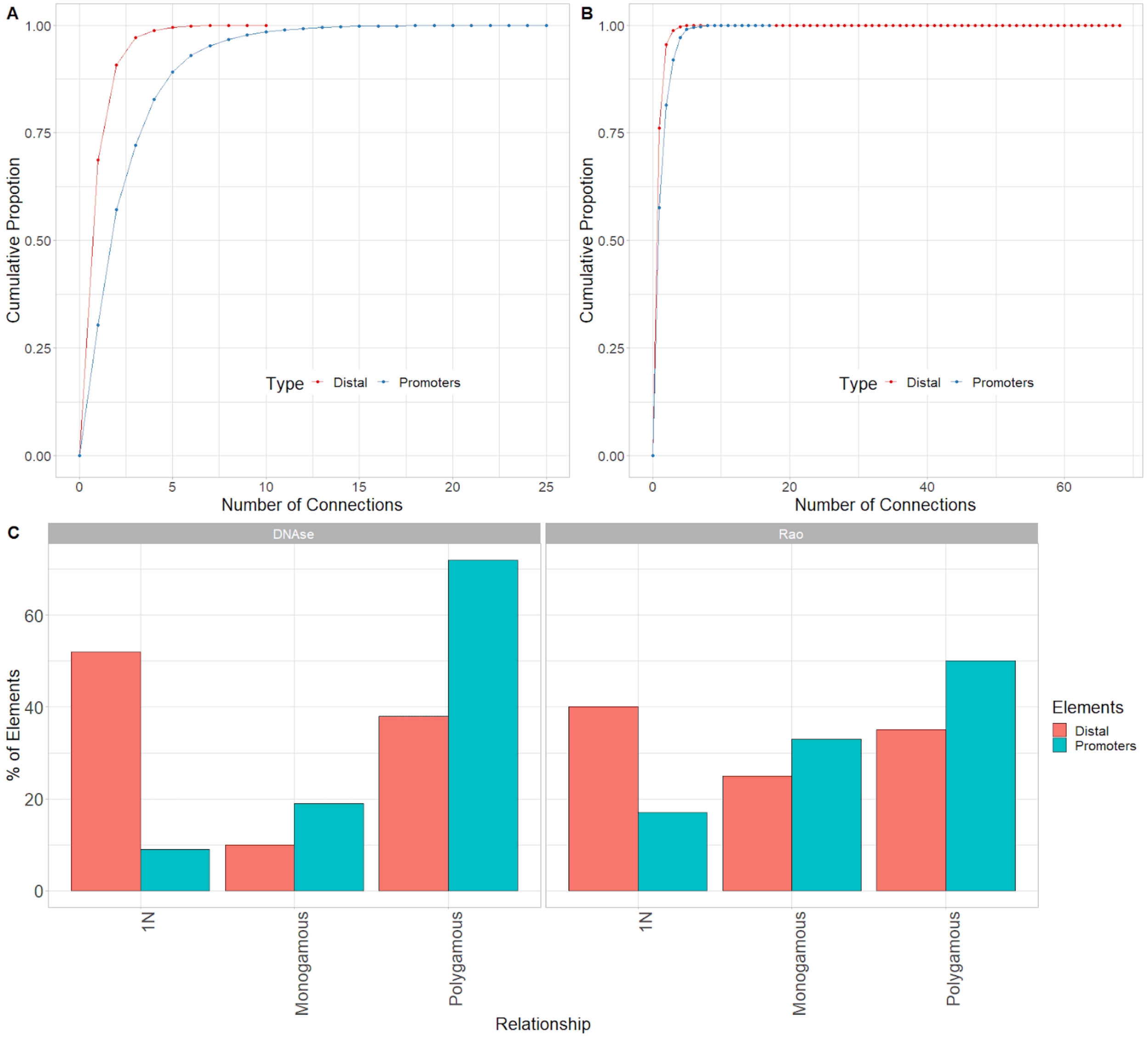


**Figure S5**

**Promoters are more connected than distal elements in DNAse-based and Rao methods**

**(A)** Cumulative proportion of the number of connections for promoters or distal elements for the DNAse method. **(B)** Cumulative proportion of the number of connections for promoters or distal elements for the Rao method. **(C)** Distribution of kind of relationship for the DNAse method (Left) Rao method (Right)**.**

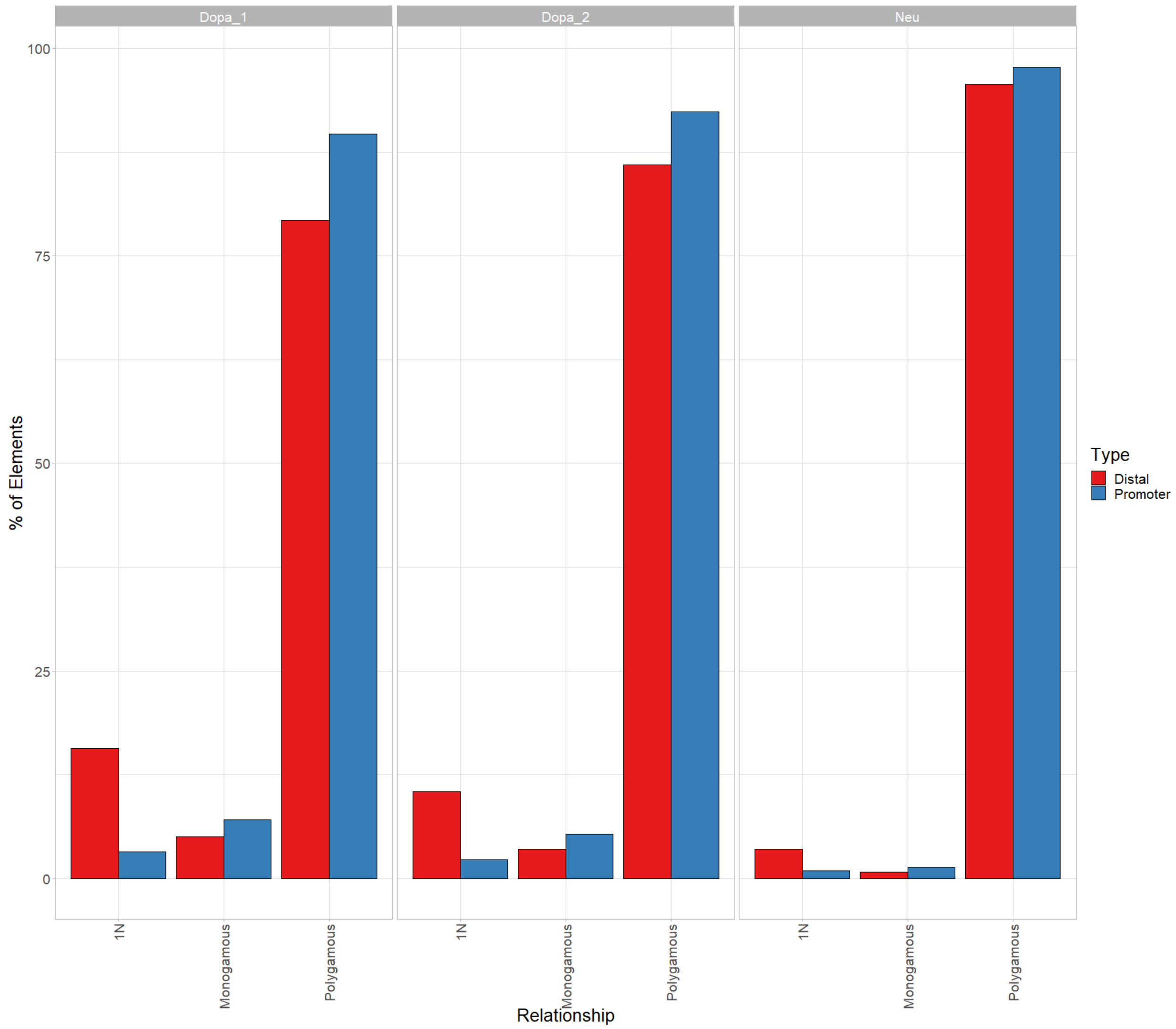


**Figure S6**

**Promoters are inside more complex relationships than distal elements in post-mortem brain tissues**

Complexity analysis for promoters and distal elements across post-mortem brain tissues.

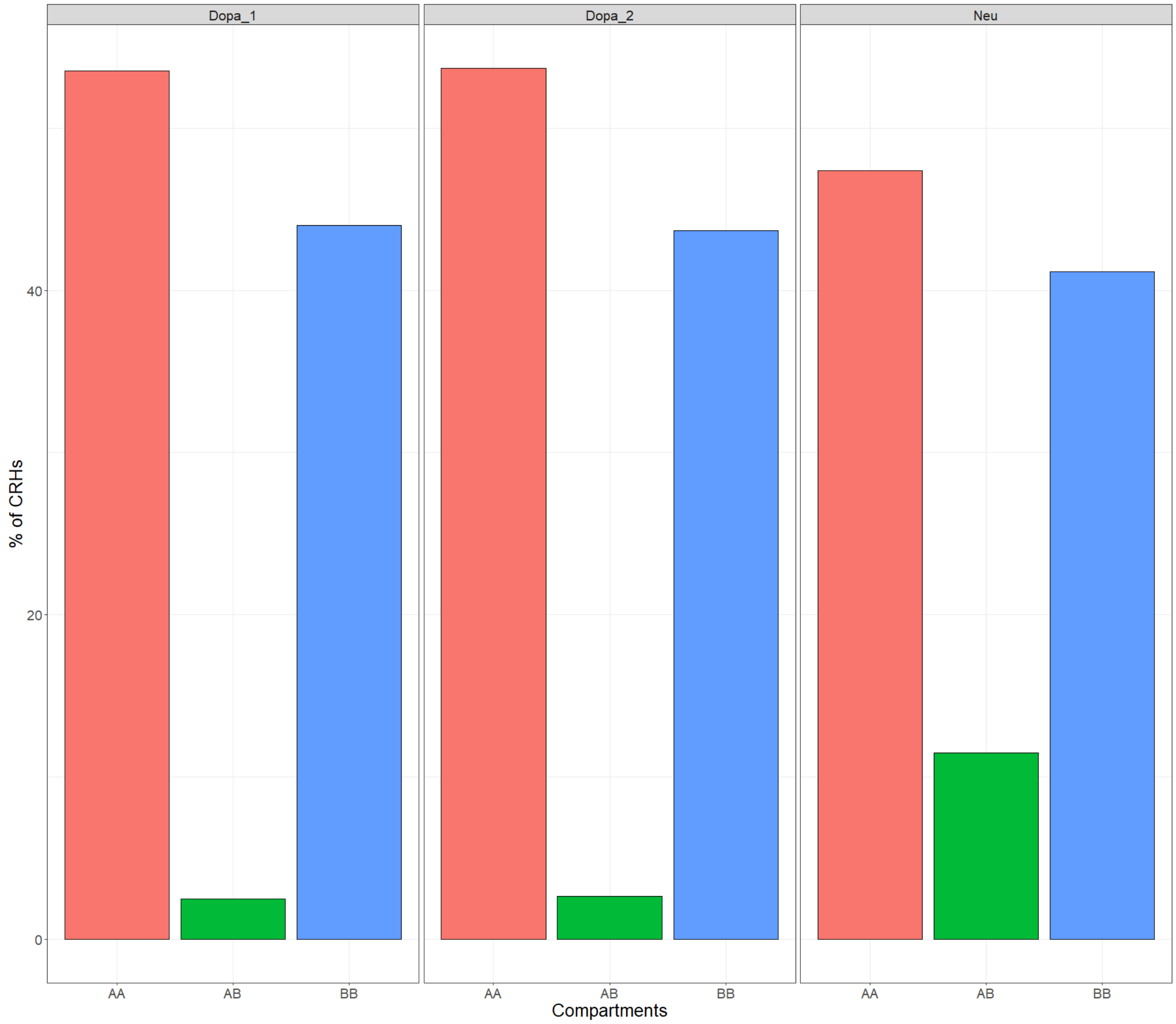


**Figure S7**

**CRHs mainly overlap active compartments across post-mortem tissues**

Distribution of the kind of compartment overlapped by CRHs across post-mortem brain tissues. Interestingly, dopaminergic neuron samples showed identical overlapping proportions.

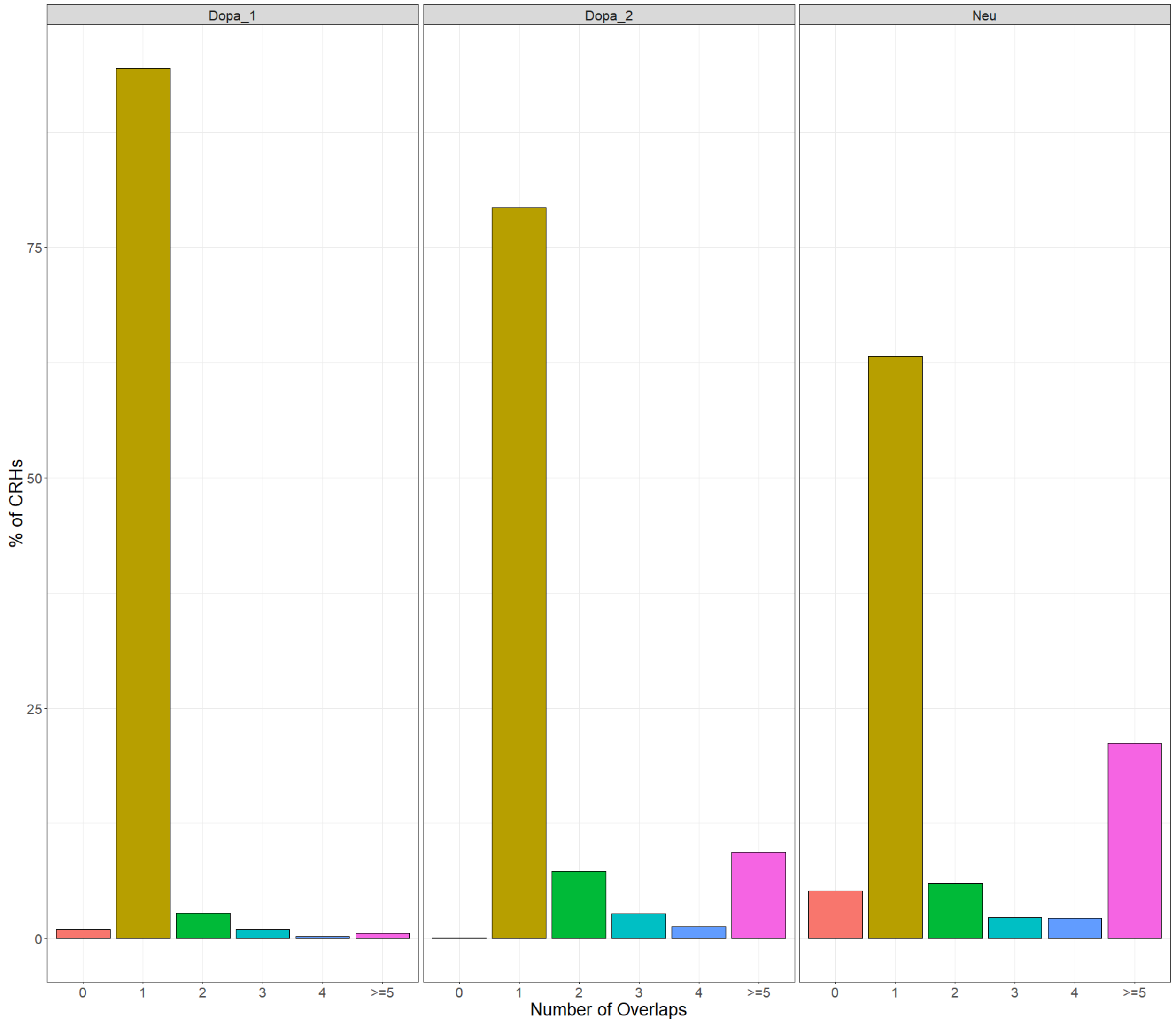


**Figure S8**

**In most cases, CRHs overlap one TAD across post-mortem brain tissues**

Distribution of the number of TADs overlapped by CRHs across post-mortem brain tissues, when TADs are detected with the directionality index.

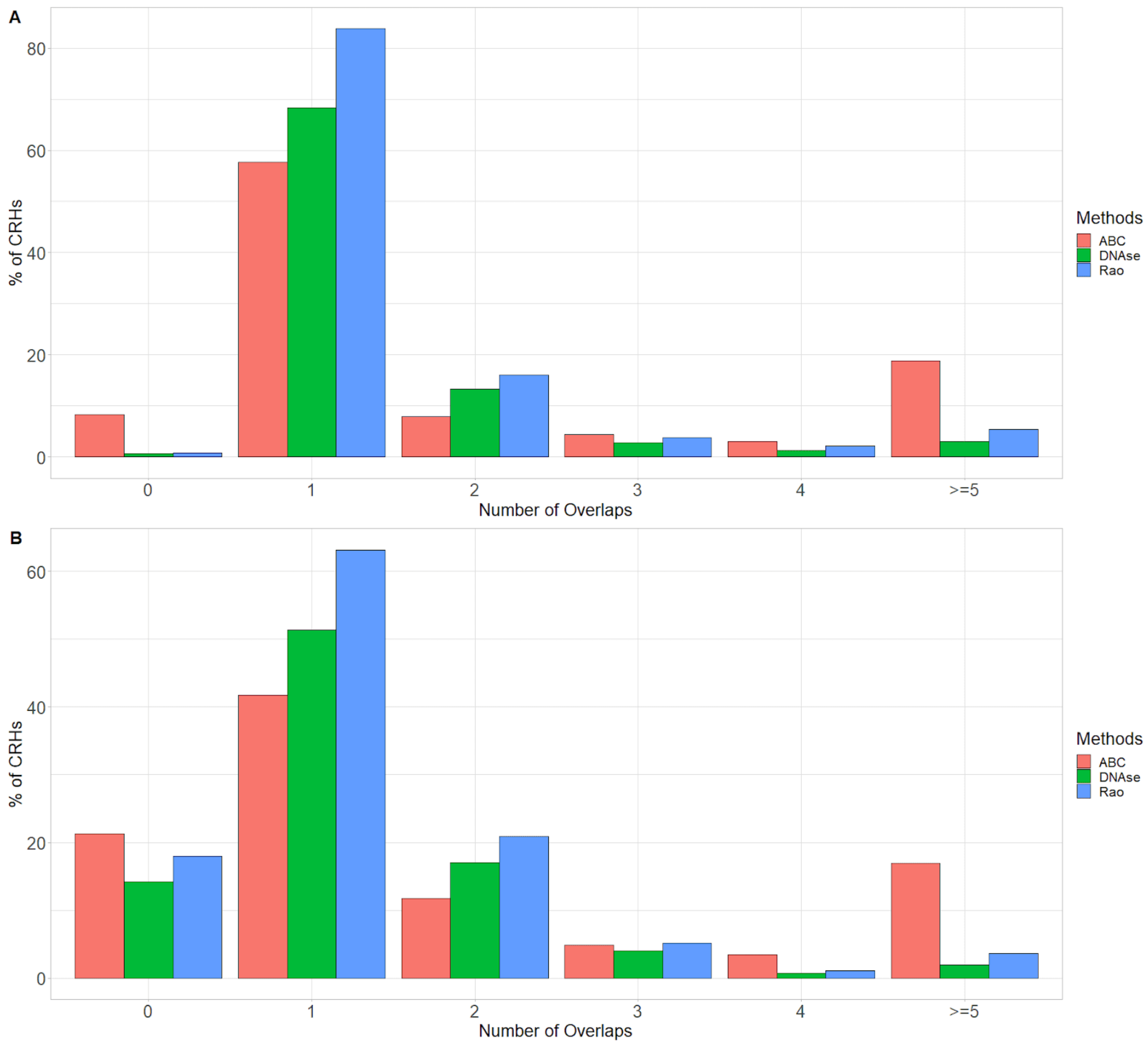


**Figure S9**

**In most cases, CRHs overlap one TAD across our methods**

**(A)** Distribution of the number of TADs overlapped by CRHs built with our different methods, when TADs are detected with the insulation score (Crane et al. 2015). **(B)** Distribution of the number of TADs overlapped by CRHs built with our different methods, when TADs are detected with the Arrowhead algorithm (Rao et al. 2014).

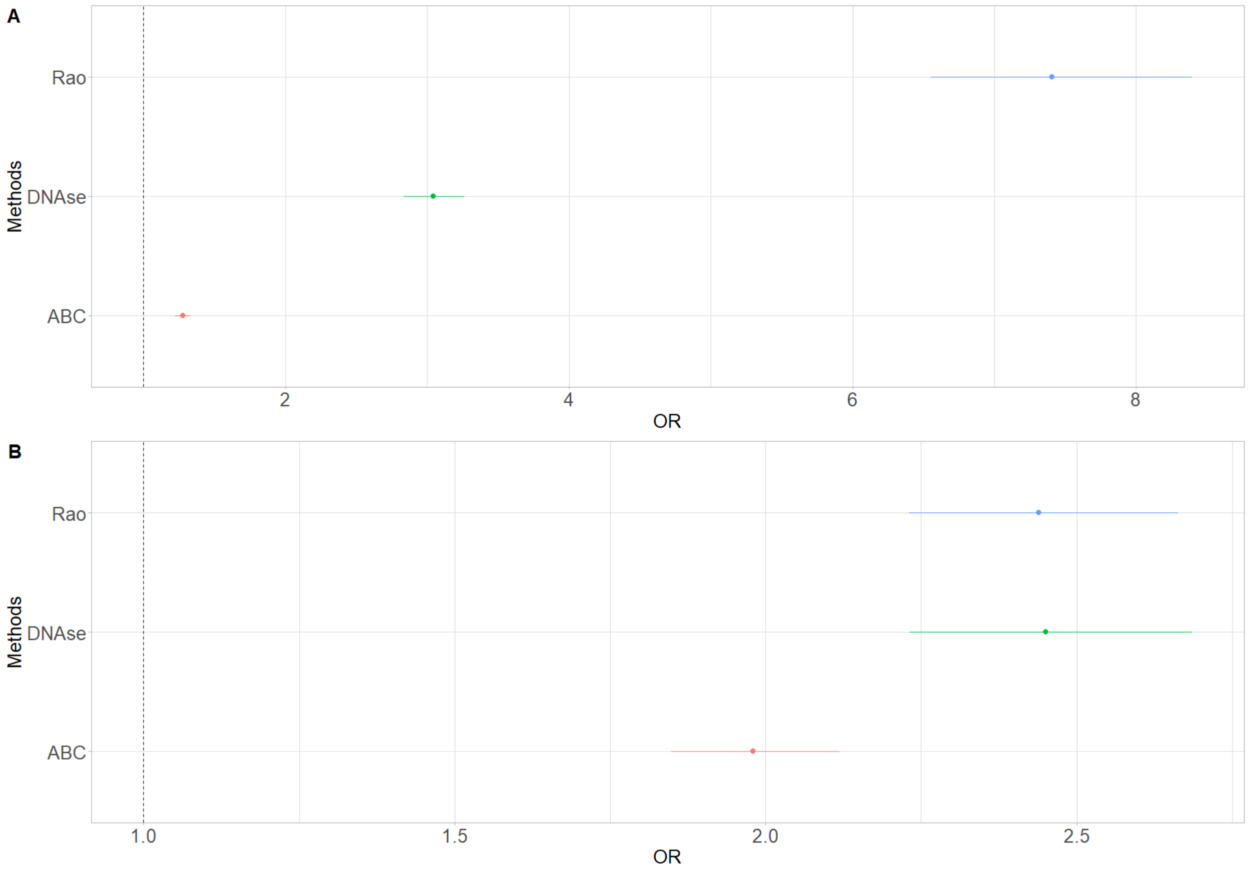


**Figure S10**

**Enrichment in FIREs for distal elements and promoters in our methods**

**(A)** Enrichment in FIREs for distal elements as measured with odds ratio (OR) and their 95% confidence interval. The dotted line represents the null value. **(B)** Enrichment in FIREs for promoters as measured with odds ratio (OR) and their 95% confidence interval. The dotted line represents the null value.

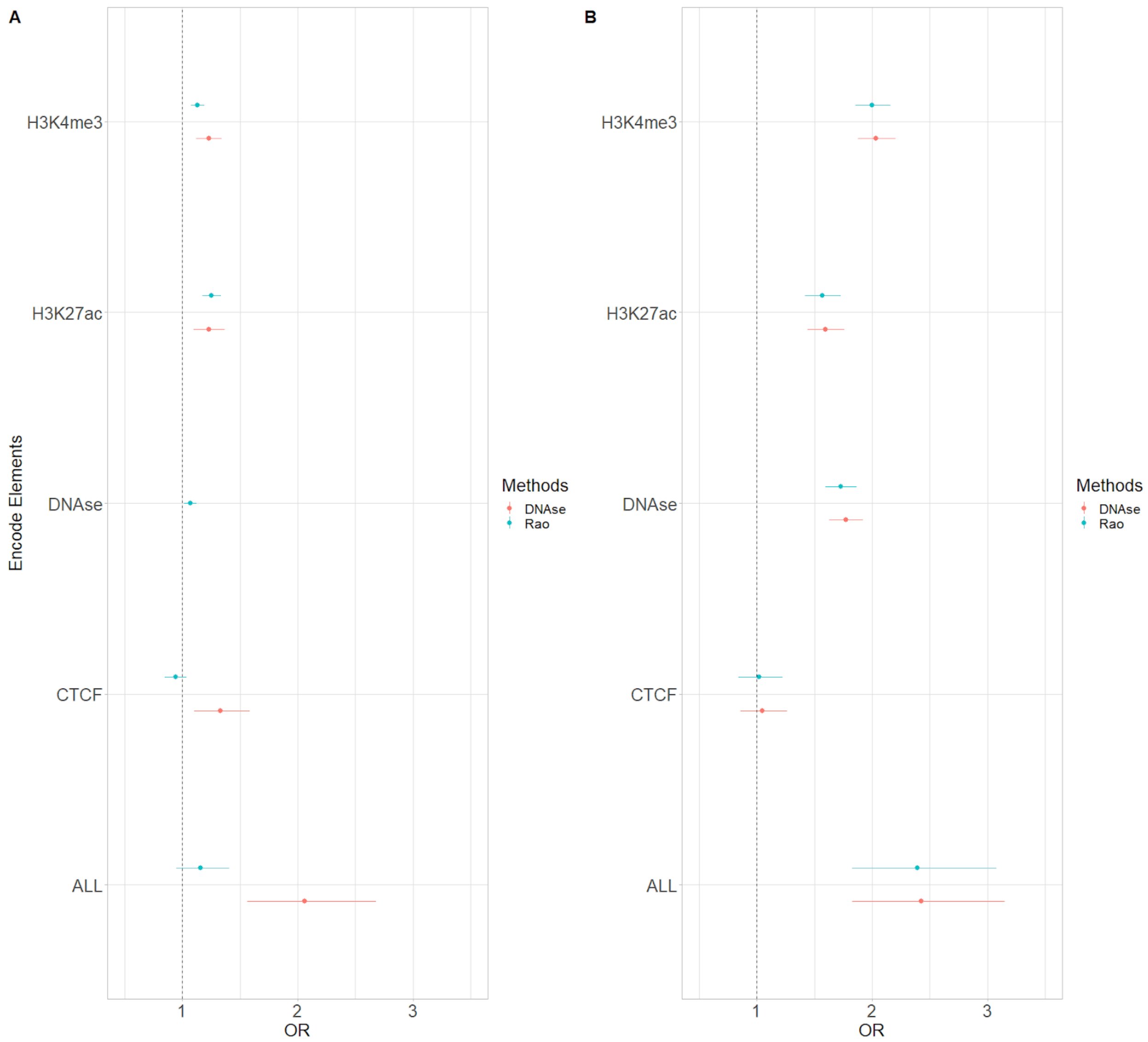


**Figure S11**

**Enrichment in Encode candidate elements for distal elements and promoters in our control methods**

**(A)** Enrichments in Encode Elements as measured with odds ratio (OR) for our control methods to build CRHs for distal elements. The dotted line represents the null value. The ALL category encompasses all significant regions for all Encode elements. Since the DNAse method is built entirely on DNAse we removed DNAse candidate regions. **(B)** Enrichments in Encode Elements as measured with odds ratio (OR) for our control methods to build CRHs for promoters. The dotted line represents the null value. The ALL category encompasses all significant regions for all Encode elements.

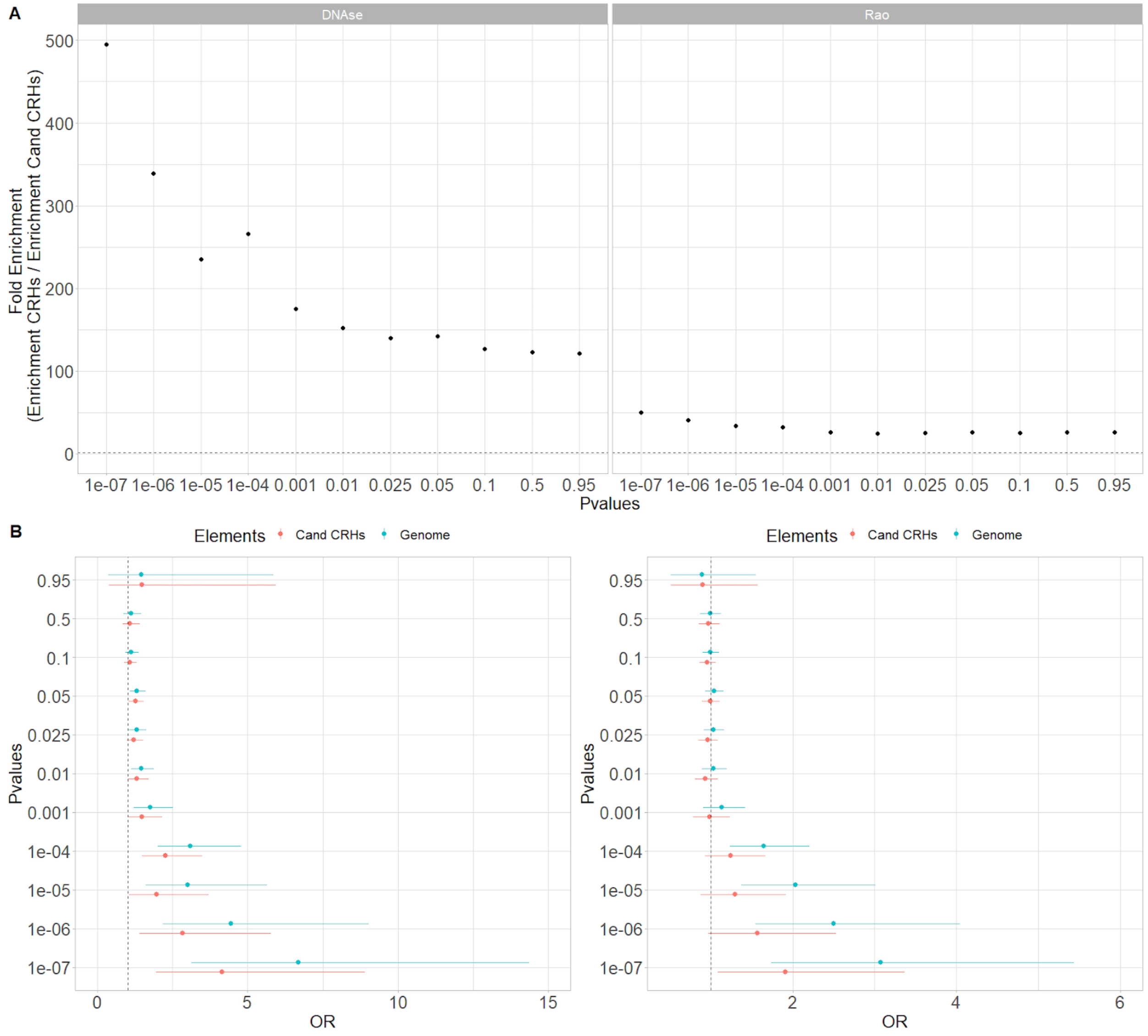


**Figure S12**

**CRHs in DNAse-based and Rao methods show strong enrichments in schizophrenia-associated SNPs**

**(A)** Fold enrichment for our control methods to build CRHs. The dotted line represents the null value. **(B)** SNP enrichments as measured with odds ratio (OR) with their 95% confidence intervals for our control methods to build CRHs. (Left) DNAse method (Right) Rao method. The dotted line represents the null value.

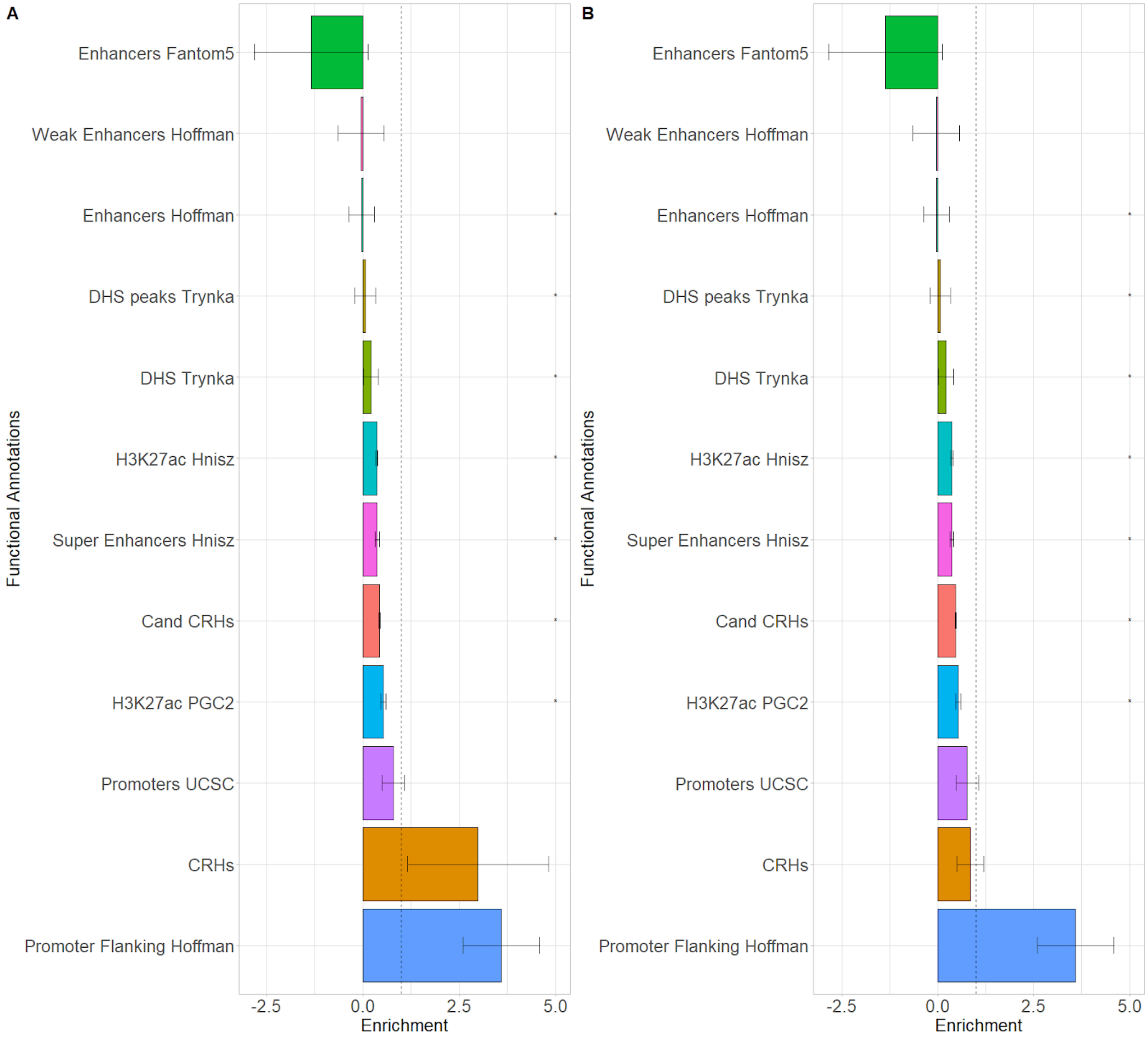


**Figure S13**

**Schizophrenia heritability enrichments for DNAse-based and Rao methods**

Schizophrenia heritability for our control methods **(A)** DNAse method **(B)** Rao method with their error bars. The dotted line represents the null.

Data Information: * represent p-value <=0.05 after Bonferroni correction.

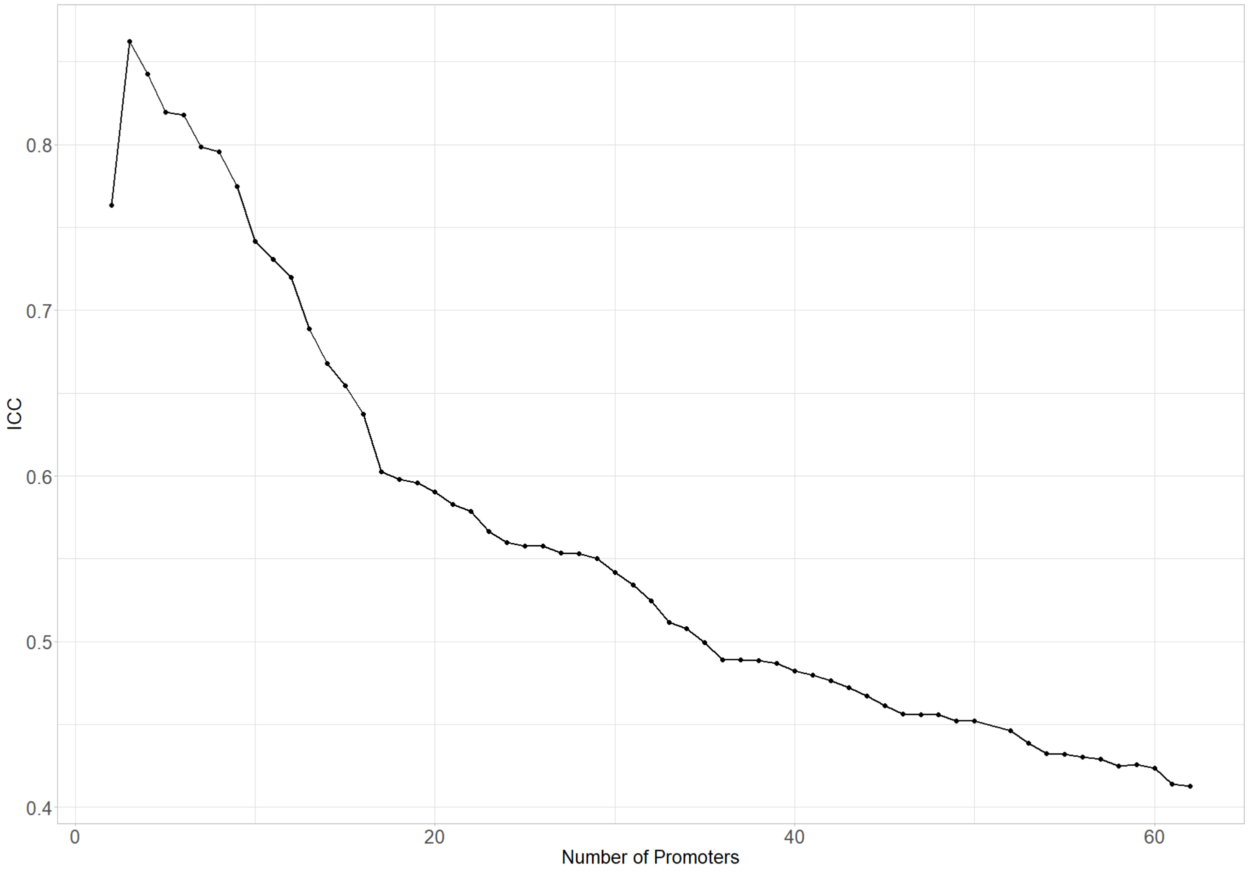


**Figure S14**

**Negative association between Intraclass correlation and the number of promoters considered within CRHs for the ABC-Score method**

Intraclass correlation (ICC) evolution of gene expression respectively of the number of promoters within the CRH.
