## Supplemental Methods for "Cis-Regulatory Hubs: a new 3D Model of Complex Disease Genetics with an Application to Schizophrenia"

### Supplementary

#### Methods

##### 3D Features

**A/B Compartments**: A/B compartments were defined at a 500 kb resolution of the contact matrix (using a 100 kb resolution had little impact on the results). The first principal component (PC) of a suitably normalized Hi-C contact matrix over a chromosome arm captures the plaid pattern of A/B compartments (Lieberman-Aiden et al., 2009). As GC content is higher in A compartments than in B compartments, the correlation of the first PC with GC content was used to orient the first PC so that positive values correspond to the A compartments and negative values to B compartments (Imakaev et al., 2012). The transformation applied to the ratio observed/expected (O/E) contact matrices was selected that 1) maximized the number of autosomal chromosome arms where the first PC had the highest correlation with GC content over the first three PCs and 2) had the highest correlation of the first PC with GC content in these chromosome arms. The transformation was selected among the following three: O/E - 1 with clipping of values below percentile 1 and above percentile 99, log (O/E) and log (O/E) with clipping of values below percentile 1 and above percentile 99. The last transformation was selected based on the above criteria, with 30/40 autosomal chromosome arms where the first PC correlated the most strongly with GC content and correlations between 0.38 and 0.87 within these chromosome arms. The A/B compartment for the remaining 10 chromosome arms (6q, 8q, 9p, 10q, 12p, 18p, 18q, 19q, 20p and 21p) were set to missing.

**TADs calling**: TADs were called using the directionality index (Dixon et al., 2012), insulation score (Crane et al., 2015) or with Arrowhead algorithm from Juicer software (Rao et al., 2014; Durand et al., 2016)

*Directionality Index* (DI) was computed as presented by Dixon et al., 2012 at 10 Kb resolution. Briefly, for each 10 Kb bins the number of upstream and downstream contacts were calculated. A bias toward upstream regions at the end of a TAD was expected and conversely, a bias toward downstream regions, at the beginning of a TAD was expected. As mentioned by Gorkin et al., 2019, the original approach to computing the DI using a 200 Kb window size was applied to capture more local features. DI values for each 10Kb bins were used to build a Hidden Markov Model and predict upstream bias, downstream bias, and no bias states, respectively. Regions switching from upstream bias to downstream bias were called topological boundaries.

*Insulation Score* (INS) was computed as presented by Crane et al., 2015. Simply, for each 10 Kb bin, the average number of contacts in 400Kb windows upstream and downstream on O/E matrices was computed. A local minimum for INS at TADs borders was expected. INS was normalized at the chromosome level to take account of differences between chromosomes. Then INS was scaled between 0 and 1, where 0 is complete insulation and 1 is no insulation respectively.

*Arrowhead* TADs were annotated using Arrowhead (Rao et al., 2014; Durand et al., 2016) at 10 Kb resolution.

**Frequently Interacting Regions** (FIREs) (Schmitt et al., 2016) was computed with FIREcaller R package (Crowley et al., 2021) at 10Kb resolution, with minor adjustments to fit our data format.

#### Functional Enrichment analysis

Genes inside CRHs were used for GO enrichment analysis at the CRH or gene level. To do so, at the CRH level the clusterProfiler R package (Wang et al., 2012) was used with the compareCluster function to perform over-representation tests.

#### Peak calling

Peak calling for ChIP-Seq data was performed with MACS2 (Zhang et al., 2008) software through
the following command:

```
macs2 callpeak \
-t bamfile \
-n alias \
-f BAM \
-g hs \
-p .1 \
--call-summits \
--outdir outputdirectory
```

#### Genome Build

All coordinates in the human genome are reported using build hg19.

#### Control Methods

As mentioned in the main analysis, results were controlled using two other methods to determine distal elements and CRHs. In doing so, a large spectrum of regulatory processes was captured. Thus, alternative CRHs were defined through the Rao and DNAse methods, described as follows.

**Rao**: Since it has been shown that significant 3D peaks are enriched in enhancers, all distal elements were defined by 3D significant peaks linking promoters (Rao et al., 2014).

Briefly, as shown by Rao et al., 2014, a significant 3D peak is defined by comparing the number of contacts in this 3D peak relative to four neighborhood regions: horizontal, vertical, lower-left and donut, respectively.

**DNAse-based**: Since it has been shown that non-coding regions are open chromatin regions, DNAse peaks were added as an additional biological layer to the 3D peaks from the Rao method. In this method, distal elements are all DNAse peaks on 10Kb 3D peaks in 3D contact with a promoter. Due to the methodology, several DNAse peaks can be observed on the same 10Kb peak. Thus, each peak was considered as an individual distal element.

Also mentioned in the main method section, we defined candidate CRHs in order to assess comparison for enrichment analysis. For the Rao method, we considered all significant 3D peaks in no contact with a promoter as candidate elements. Based on the same rationale, DNAse candidate elements are all DNAse peaks in no 3D contact with a promoter.

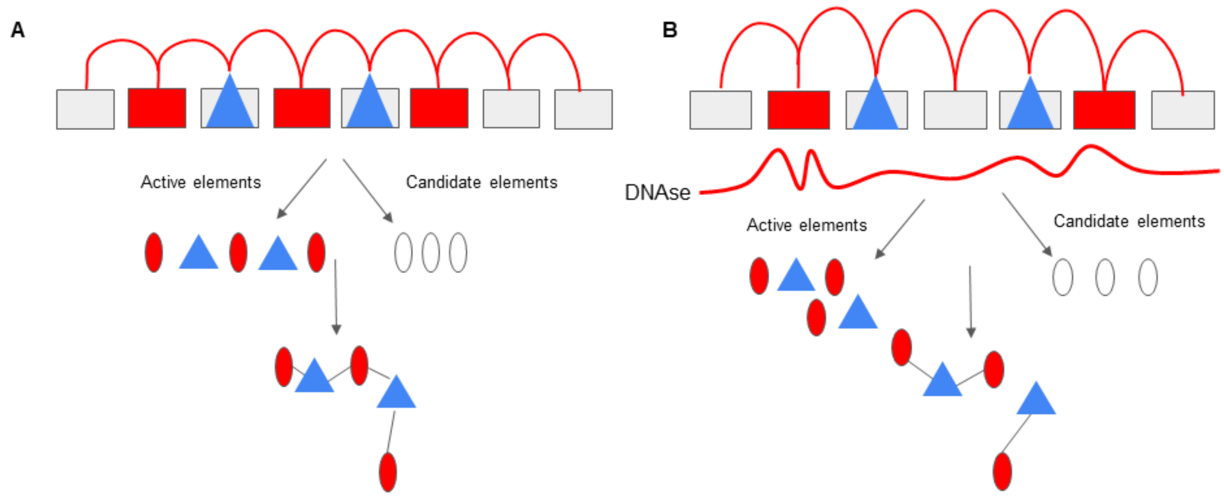


**Figure Supplemental Methods 1**

**(A)** Rao-Based method methodology and CRH building. **(B)** DNAse-based method methodology and CRH building

Data Information: Promoters are represented by blue triangles, distal regulatory elements by red circles and, candidate elements by white circles.

### HiC Mapping, Filtering, and Normalization for post-mortem brains

Raw Hi-C sequence fastq files for postmortem dopaminergic neuronal nuclei (NeuN+/Nurr1+) and the general neuronal (NeuN+) populations were obtained from the PsychENCODE Synapse platform. We referred in supplementary figures to Dopa_1 and Dopa_2 for postmortem dopaminergic neuronal nuclei samples and Neu for the general neuronal sample, respectively. They were mapped to the human genome sequence (hg19) in 10 kb bins using distiller (https://github.com/mirnylab/distiller-nf). Genome-wide iterative correction (i.e., KR normalization) was performed using cooler (https://github.com/mirnylab/cooler). We used the hicConvertFormat tool from the HiCExplorer package (https://hicexplorer.readthedocs.io) to convert .cool files into ginteractions files, which after preprocessing were read by the Juicer toolbox and converted to .hic files.

#### Acknowledgments

The PsychEncode project is supported by: U01MH103392, U01MH103365, U01MH103346, U01MH103340, U01MH103339, R21MH109956, R21MH105881, R21MH105853, R21MH103877, R21MH102791, R01MH111721, R01MH110928, R01MH110927, R01MH110926, R01MH110921, R01MH110920, R01MH110905, R01MH109715, R01MH109677, R01MH105898, R01MH105898, R01MH094714, P50MH106934, U01MH116488, U01MH116487, U01MH116492, U01MH116489, U01MH116438, U01MH116441, U01MH116442, R01MH114911, R01MH114899, R01MH114901, R01MH117293, R01MH117291, R01MH117292 awarded to: Schahram Akbarian (Icahn School of Medicine at Mount Sinai), Gregory Crawford (Duke University), Stella Dracheva (Icahn School of Medicine at Mount Sinai), Peggy Farnham (University of Southern California), Mark Gerstein (Yale University), Daniel Geschwind (University of California, Los Angeles), Fernando Goes (Johns Hopkins University), Thomas M. Hyde (Lieber Institute for Brain Development), Andrew Jaffe (Lieber Institute for Brain Development), James A. Knowles (University of Southern California), Chunyu Liu (SUNY Upstate Medical University), Dalila Pinto (Icahn School of Medicine at Mount Sinai), Panos Roussos (Icahn School of Medicine at Mount Sinai), Stephan Sanders (University of California, San Francisco), Nenad Sestan (Yale University), Pamela Sklar (Icahn School of Medicine at Mount Sinai), Matthew State (University of California, San Francisco), Patrick Sullivan (University of North Carolina), Flora Vaccarino (Yale University), Daniel Weinberger (Lieber Institute for Brain Development), Sherman Weissman (Yale University), Kevin White (University of Chicago),

Jeremy Willsey (University of California, San Francisco), and Peter Zandi (Johns Hopkins University).
